# Assessing deep learning algorithms in *cis*-regulatory motif finding based on genomic sequencing data

**DOI:** 10.1101/2020.11.30.403261

**Authors:** Yan Wang, Shuangquan Zhang, Anjun Ma, Cankun Wang, Zhenyu Wu, Dong Xu, Qin Ma

## Abstract

*Cis*-regulatory motif finding is a crucial step in the detection of gene regulatory mechanisms using genomic data. Deep learning (**DL**) models have been utilized to denovoly identify motifs, and have been proven to outperform traditional methods. By 2020, twenty DL models have been developed to identify DNA and RNA motifs with diverse framework designs and implementation styles. Hence, it is beneficial to systematically compare their performances, which can facilitate researchers in selecting the appropriate tools for their motif analyses. Here, we carried out an in-depth assessment of the 20 models utilizing 1,043 genomic sequencing datasets, including 690 ENCODE ChIP-Seq, 126 cancer ChIP-Seq, 172 single-cell cleavages under targets and release using a nuclease, and 55 RNA CLIP-Seq. Four metrics were designed and investigated, including the accuracy of motif finding, the performance of DNA/RNA sequence classification, algorithm scalability, and tool usability. The assessment results demonstrated the high complementarity of the existing models, and it was determined that the most suitable model should primarily depend on the data size and type as well as the model outputs. A webserver was developed to allow efficient access of the identified motifs and effective utilization of high-performing DL models.

## 1. Introduction

Transcription factors (**TF**s) possess unique gene expressions by binding to specific DNA or RNA sequences, named TF binding sites (**TFBS**s), and the aligned TFBSs are referred to as *cis*-regulatory motifs (*motifs* for short). It is well known that TFs are closely related to disease occurrence and progression, and motif finding and analyses can greatly benefit research of human health^1^. Substantial computational efforts have been carried out to identify TFBS specificity and predict motif patterns from the Chromatin Immunoprecipitation Sequencing (**ChIP-Seq**) data and crosslinking-immunoprecipitation and high-throughput sequencing (**CLIP-Seq**) data, which provides genome-wide interactions between TFs and DNA sequences. Meanwhile, the emergence of Cleavage Under Target & Release Using Nuclease (**CUT&RUN**) allowed for the identification of low-signal genomic features that were undetected by ChIP-Seq, providing more accurate quantifications of TF-DNA interactions and identifying motif patterns in single cells^2,3^. However, considering the complexity of TF binding mechanisms, advanced computational approaches are needed to derive biologically meaningful insights from the above large-scale genomic sequencing data.

Moreover, the increasing need in identifying the heterogenous TF-DNA interactions at the single-cell level, using the CUT&RUN technique, greatly increases the data sparsity and dimensionality, leading to a strong requirement of more efficient and powerful tools.

Deep learning (**DL**) technology is an unprecedented method exhibiting tremendous progress on motif finding and TFBSs identification from the ChIP-Seq and CLIP-Seq data^4^. Different DL models have been developed for ChIP-Seq data, including convolutional neural networks (**CNN**)^5^, recurrent neural networks (**RNN**)^6^, deep belief network (**DBN**)^7^, and graph convolutional neural network (**GCNN**)^8^, resulting in more than 20 computational tools (e.g., DESSO, DeepBind, etc.). Given the diversity of DL models in motif prediction, it is important to quantitively assess their performances and robustness across a large number of different datasets. Furthermore, how these existing tools can be appropriately applied to cancer-related data and single-cell data remains unknown. Without a clear assessment, new researchers in this field will struggle in choosing the most appropriate tool for their specific studies related to gene regulation.

To address this problem, we benchmarked the existing 20 DL tools^9–28^ (**Supplementary Table S1**) using 690 ENCODE DNA ChIP-Seq datasets (covering 91 TFs in 161 cell lines) and 55 RNA CLIP-Seq datasets (**Fig. 1A-B** and (**Supplementary Table S2**) ^28,23^. The 20 tools are separated into two categories: CNN network and hybrid network (a combination of CNN and RNN/DBN), and further summarized into five algorithmic architectures (**Fig. 1C**). To benchmark the tools, we evaluated the overall performance for DNA/RNA sequence classification, motif finding, algorithm scalability, and tool usability. Furthermore, we applied these tools on 126 cancer-related ChIP-Seq datasets and 172 single-cell CUT&RUN datasets^3^ to evaluate the capability of these DL models in elucidating shared and specific motifs among cancer types and cell groups (**Supplementary Table S2**). We summarize the performance of the 20 tools based on our assessment results, with the purpose of assisting researchers in deciding the appropriate tools for their motif analysis studies. To leverage the reproducibility of our work and also lessen the burden of using these DL tools, we developed a webserver to allow the efficient access of the identified motifs and effective utilization of high-performing DL models.

**Fig. 1.**
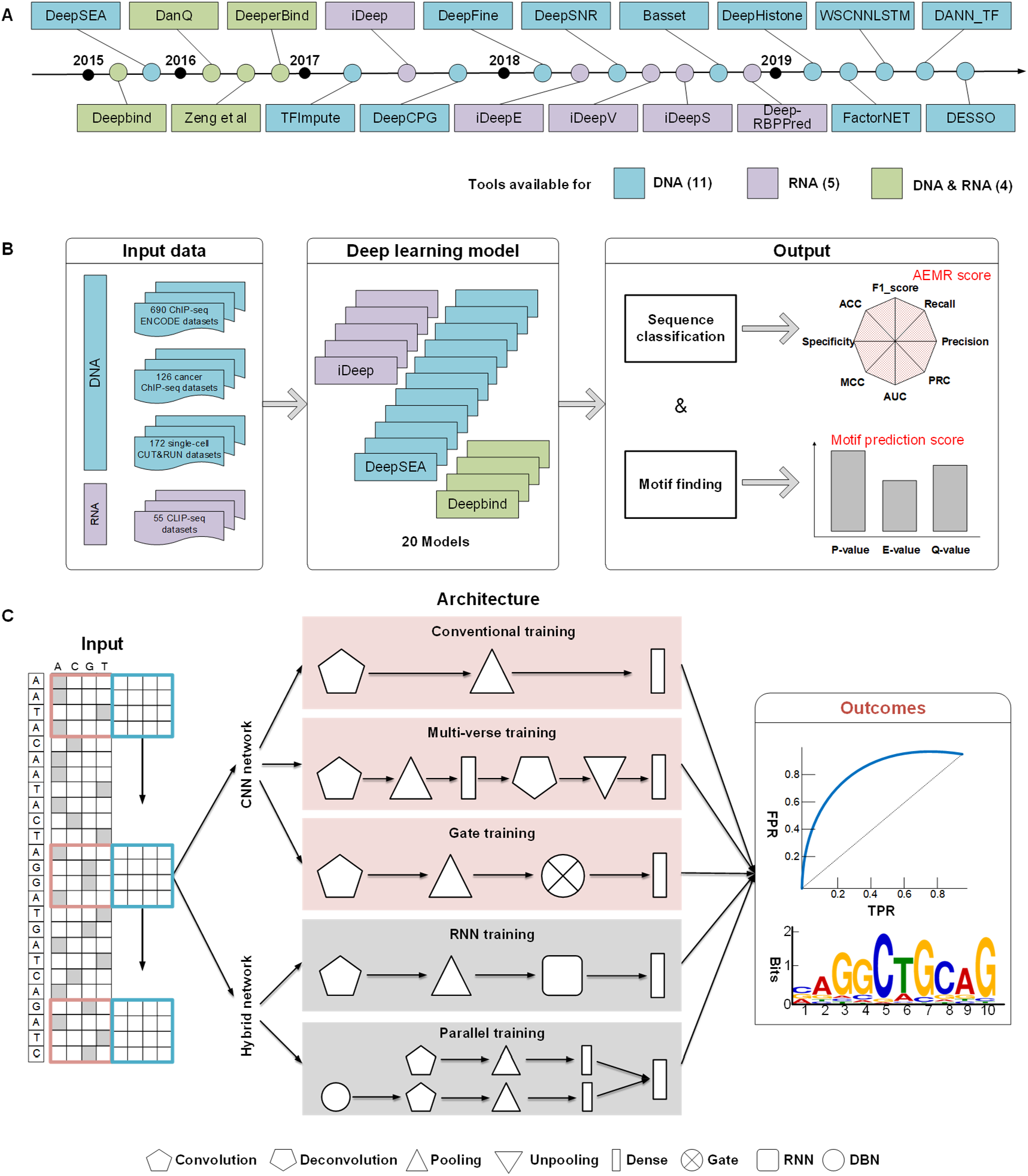
Overview of the DL tools categorization and evaluation pipeline. (*A*) The selected 20 DL tools are arranged along their publication timeline. (*B*) Schematic overview of the evaluation pipeline. We defined two scores to evaluate the quality of sequence classification and motif finding of each tool. The area of eight matrices radar (AEMR) score assesses the sequence classification ability based on F1_score, recall, precision, precision-recall curve (PRC), area under the curve (AUC), Matthews correlation coefficient (MCC), specificity, and accuracy (ACC) between predicted classification labels and ChIP-Seq peak labels. A motif prediction score assesses the alignment significance of predicted motifs to documented TFBSs based on *P*-values, *E*-values, and *Q*-values (see the Evaluation workflow in the Methods section). (*C*) Detailed ChIP-Seq data input and DL model classification based on CNN or hybrid networks and different framework architectures. Outcomes including both predicted sequence labels and identified motif patterns.

## 2. Results

### 2.1. Comprehensive benchmarking of 20 tools using 745 datasets and four evaluation metrics

We first benchmarked the tool performance in analyzing DNA and RNA sequences, respectively (**Fig. 2**). The 20 tools were grouped into CNN network-based and hybrid network-based according to the architecture illustrated in **Fig. 1C**. To systematically compare the 20 tools, we designed four metrics to assess their performances from multiple aspects: (*i*) an area of eight metrics radar (**AEMR**) score defines the refined accuracy of DNA/RNA sequence classification, (*ii*) a motif prediction score to assess the significance and accuracy of identified motifs, (*iii*) algorithm scalability of running time performed on different data scales, and (*iv*) usability of the tool in terms of the tutorial, updates, code quality, etc. (**Supplementary Table S3** and **Fig. S1**).

**Fig. 2.**
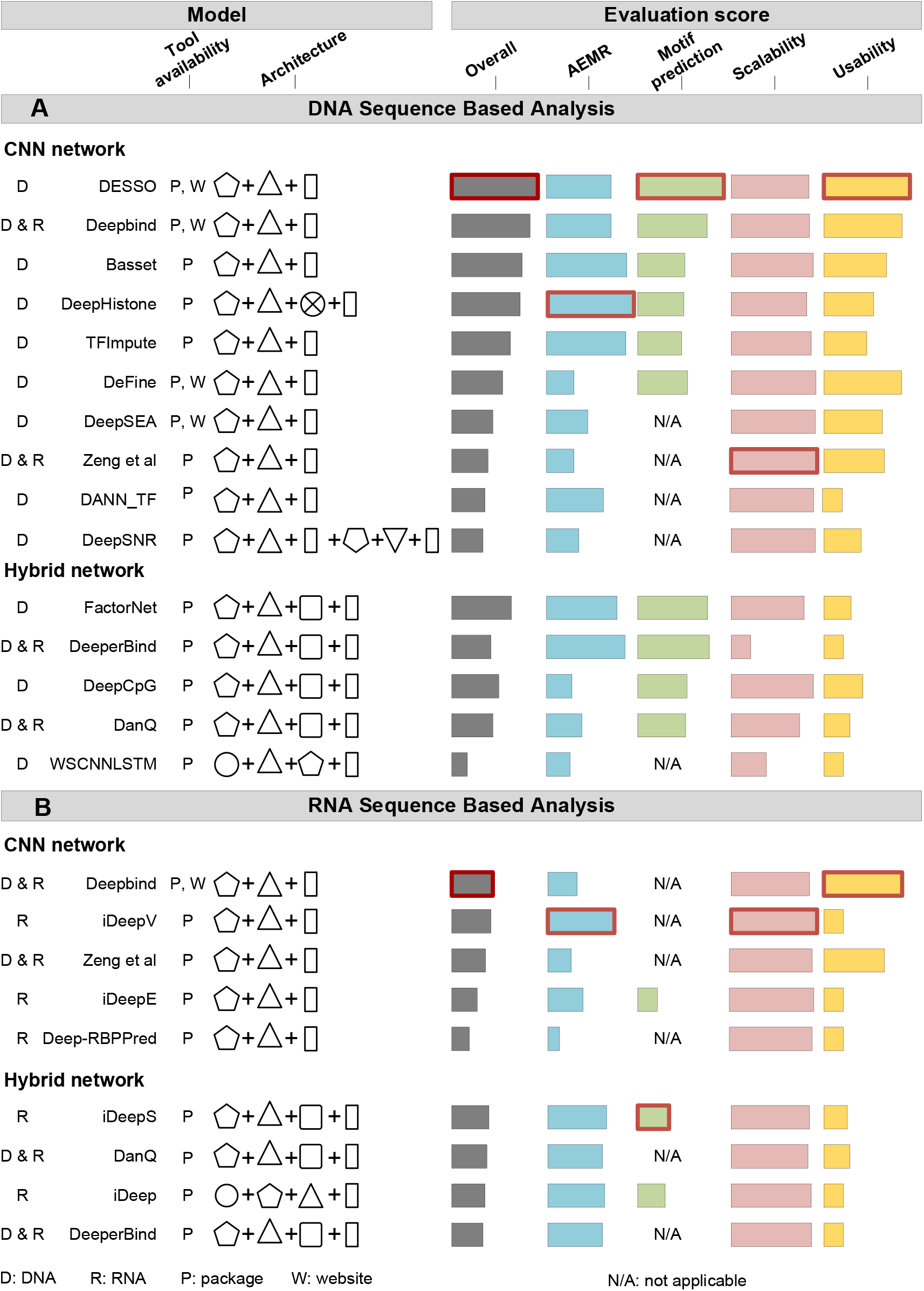
Illustration of the 20 tools with overall evaluation results. (A) For DNA sequence based analysis, tools were separated by DL models (CNN network and hybrid network). Tool name, tool availability (package/website), and summarized architecture model were given for each tool. In each comparative group, tools were ranked by their overall score (grey) from high to low. Four evaluation scores were shown: AEMR (blue), motif prediction score (green), algorithm scalability (pink), and tool usability (yellow). The highest score for each evaluation score is highlighted in red. (B) For RNA sequence based analysis, the same columns and labels were used as described in A.

Based on all the above four metrics, an overall score was calculated to evaluate the 20 DL tools, and then was used, to rank each tool in the decreasing order of its value. As shown in **Fig. 2**, tools’ performance differed the most in the AEMR score, motif prediction score, and tool usability, while the algorithm scalability scores were moderately varied. DESSO achieves the highest overall score for DNA sequence based analysis than all the other DL tools (**Fig. 2A**), and DeepBind is the best tool for RNA sequence based analysis (**Fig. 2B**), also the second-best tool for DNA (**Supplementary Table S4)**. Interestingly, our results showcased that, most CNN network tools are better than hybrid network tools for DNA sequence based analysis, while worse for RNA sequence based analysis (**Fig. 2** and **Supplementary Fig. S2**). We reasoned that such results may due to insufficient data training of RNA motifs and more noises in RNA CLIP-Seq data, comparing to DNA ChIP-Seq data.

The AEMR and motif prediction scores are the most important scores that directly assess the output results of the DL models and determine the overall score. For DNA models (the DL models in **Fig. 2A**), DeepHistone showed the best performance in terms of the AEMR score (**Fig. 3A**), and DESSO is the best tool to identify the most abundant and documented motif patterns (**Fig. 3B**). Five out of 15 tools available for DNA sequence based analysis lack the ability to predict motifs from ChIP-Seq data, leading to lower overall scores than the others (**Fig. 3B**). These five tools were excluded from the evaluation of cancer data and single-cell CUT&RUN data analysis in the following sections.

**Figure 3.**
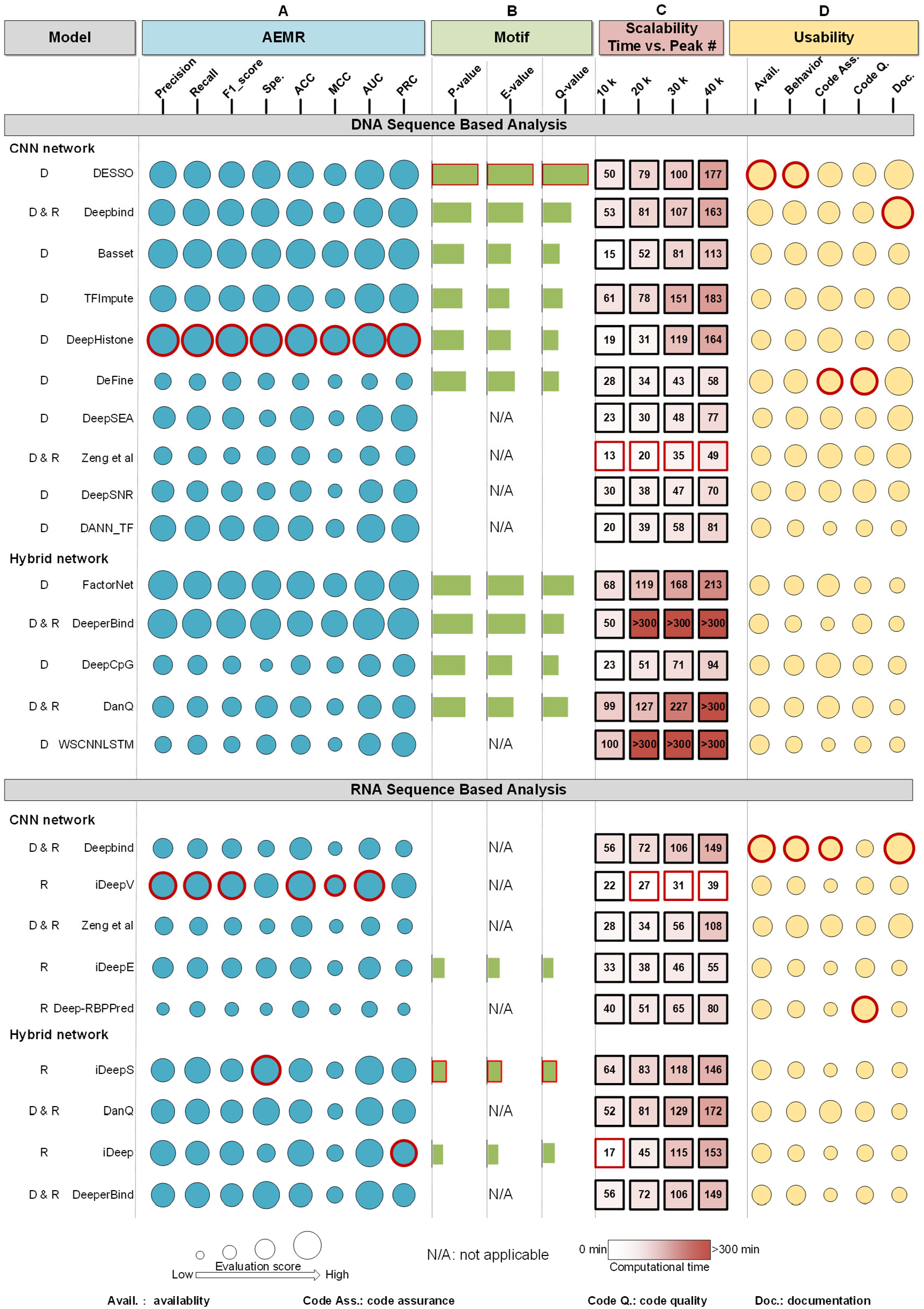
Detailed results for the four main evaluation criteria: AEMR score, motif prediction score, scalability, and usability. Note that the run times strongly depend on the implementation, hardware, dataset dimensions, and parameter settings. Therefore, run time occupies a smaller proportion in our evaluation. The best-performed tool for each sub-metrics was highlighted by red, per DNA, and RNA model.

Running times were recorded by applying each tool for different peak numbers from 10-40 thousand. All CNN network models showed higher scalability scores than hybrid network models, with an exception of DeepCpG which runs faster and steadily for large dataset analysis (**Fig. 3C**). Tools that only perform DNA sequence classification have an obvious advantage in achieving better scalability scores. To benchmark the tool usability, we considered five criteria, including tool availability, update behavior, code assurance, code quality, and documentation (**Fig. 4D** and **Methods**). DESSO is considered to be the easiest accessible tool with a user-friendly webserver and detailed documentations, while most tools with only packages available usually need more effort in installation and environment settings. DeepBind is also objected to being in good maintenance and outperformed tutorial documentation, especially for RNA sequence based analysis. For RNA models (the DL models in **Fig. 2B**), iDeepV and iDeepS the best tool for RNA sequence classification and RNA motif identification, respectively.

**Fig. 4.**
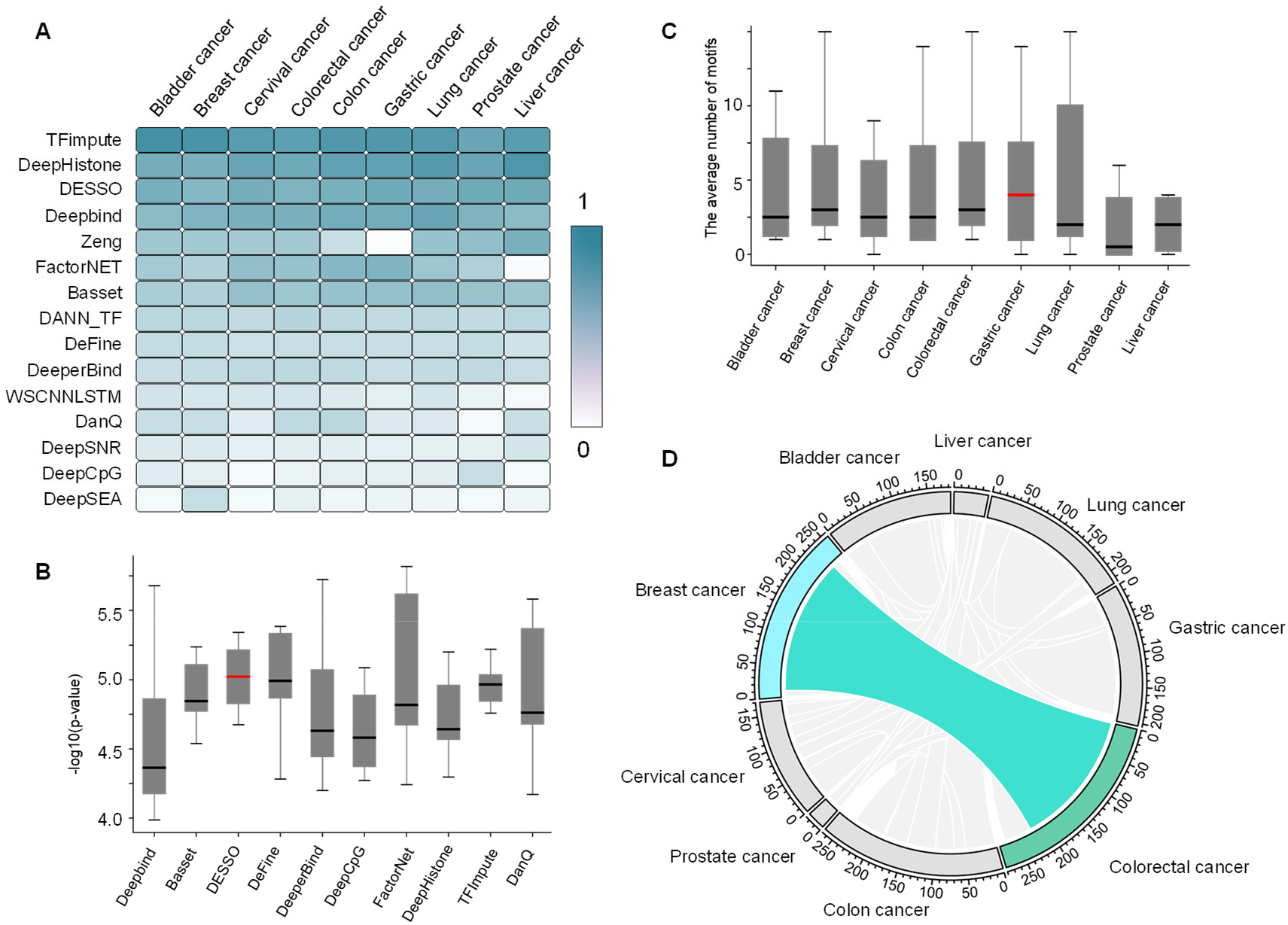
Analysis of 15 DNA models in nine cancer types. (A) AEMR scores for the DNA models across the nine cancer types. (B) Box plot of motif enrichment p-value (details in Method section) of ten DL models with respect to breast cancer. The horizontal red line indicates the highest median value on the y-axis. (C) The average number of motifs for each cancer type. The horizontal red line indicates the highest median value on the y-axis. (D) The shared motifs between the nine different cancer types.

### 2.2 DNA models found significant motifs on nine cancer types

The TF-DNA binding mechanisms are closely related to cancer occurrence and progression ^1^, and we are interested in how these DL tools (DNA models) can perform analysis on and derive new insights from cancer-related ChIP-Seq data. The five DL tools without the motif prediction function were excluded, and the remained ten tools were used to analyze 126 ChIP-Seq datasets from nine cancer types (downloaded from Cistrome Data Browser as of 04/08/2020). An overview result showcased sequence classification was very variable across tools, and TFimpute had the highest performance than the other tools (**Fig. 4A** and **Supplementary Fig. S3** and **Table S5**). DeepHiston and DeepBind performed better on liver cancer and lung cancer data, respectively. It is noteworthy that, the rank of the ten tools on cancer data, in terms of classification performance, almost supports the same performance rank on benchmark data (**Fig. 3**), indicating the robustness of our benchmark design. On the other hand, minimum diversities of classification performances existed across cancer types per tool, showing that fundamentally these tools performed steadily on different datasets.

For all the identified motifs from the cancer datasets, we performed clustering using similarity scores from TOMTOM for them, and the most significant motif in terms of its *P*-value in each cluster was defined as the representative motif. Eventually, 341 representative motifs were identified from all the 126 cancer datasets for the following analyses. Interestingly, for an individual cancer type, significant diversities and variances of motif prediction performance were observed among different tools, in which TFimpute had the lowest variance and most robustness and DESSO showed the highest mean performance (**Fig. 4B**). The average number of predicted motifs across nine cancer types varied, and apparently, there was a lower number of predicted motifs in prostate cancer and liver cancer, compared with the other seven cancer types. This may due to the less abundant known motifs identified for these two cancer types than other types (**Fig. 4C**). More efforts may be needed to select a suitable tool (or a combination of multiple tools) for motif prediction than sequence classification in different cancer types.

To understand the overlap and uniqueness status of the identified 341 motifs, we used breast cancer as an example to discover its shared motifs with other cancer types. Of all 257 motifs identified in breast cancer, 220 of them can also be identified in colorectal cancer data. For example, STAT3^29,30^, FOXO3^31,32^, and FOXP1^33,34^, have been proven to play similar roles in breast and colorectal cancer. Overall, only 63 motifs are unique to its corresponding cancer type, and all the other motifs are highly shared among different cancer types (**Fig. 4D**). Those uniquely identified motifs might be the key to determine gene signatures that play essential roles in the occurrence and development of the specific cancer type. For example, ETV1 uniquely identified in breast cancer was found to have higher expression level compared to normal tissues, while the overexpression of COP1 led to a significant reduction of ETV1 and suppressed cell migration and invasion^35^. Motif related to KLF4 identified in colorectal cancer is a zinc finger TF, which has been confirmed to be a tumor suppressor gene^36^.

### 2.3 Motif analysis on single-cell CUT&RUN data reveals potential TF co-regulatory mechanisms

The 10 DL tools used above were also applied to 172 mouse embryonic stem cells of single-cell CUT&RUN sequencing (including 120 cells for CTCF, 26 cells for SOX2, and 26 cells for NANOG) to identify TF co-regulatory patterns of specific genes ^3^ (**Supplementary Fig. S4-S5**). We identified 118, 124, and 205 motifs from SOX2, NANOG, and CTCF data, respectively, which were significantly matched to documented TFBSs in the HOCOMOCO database^37^. Using the same strategy of motif clustering in Section 2.2, 240 representative motifs were remained in the 172 cells for the following analysis (**Supplementary Table S6**), including 65, 17, and 13 motifs that are uniquely identified in CTCF, SOX2, and NANOG data, respectively. Additionally, 62 motifs shared among SOX2, NANOG, and CTCF (Group 1), 44 motifs shared between NANOG and CTCF (Group 2), 34 motifs shared between SOX2 and CTCF (Group 3), and five motifs shared between SOX2 and NANOG (Group 4) (**Fig. 5A-B** and **Supplementary Table S7**).

**Fig. 5.**
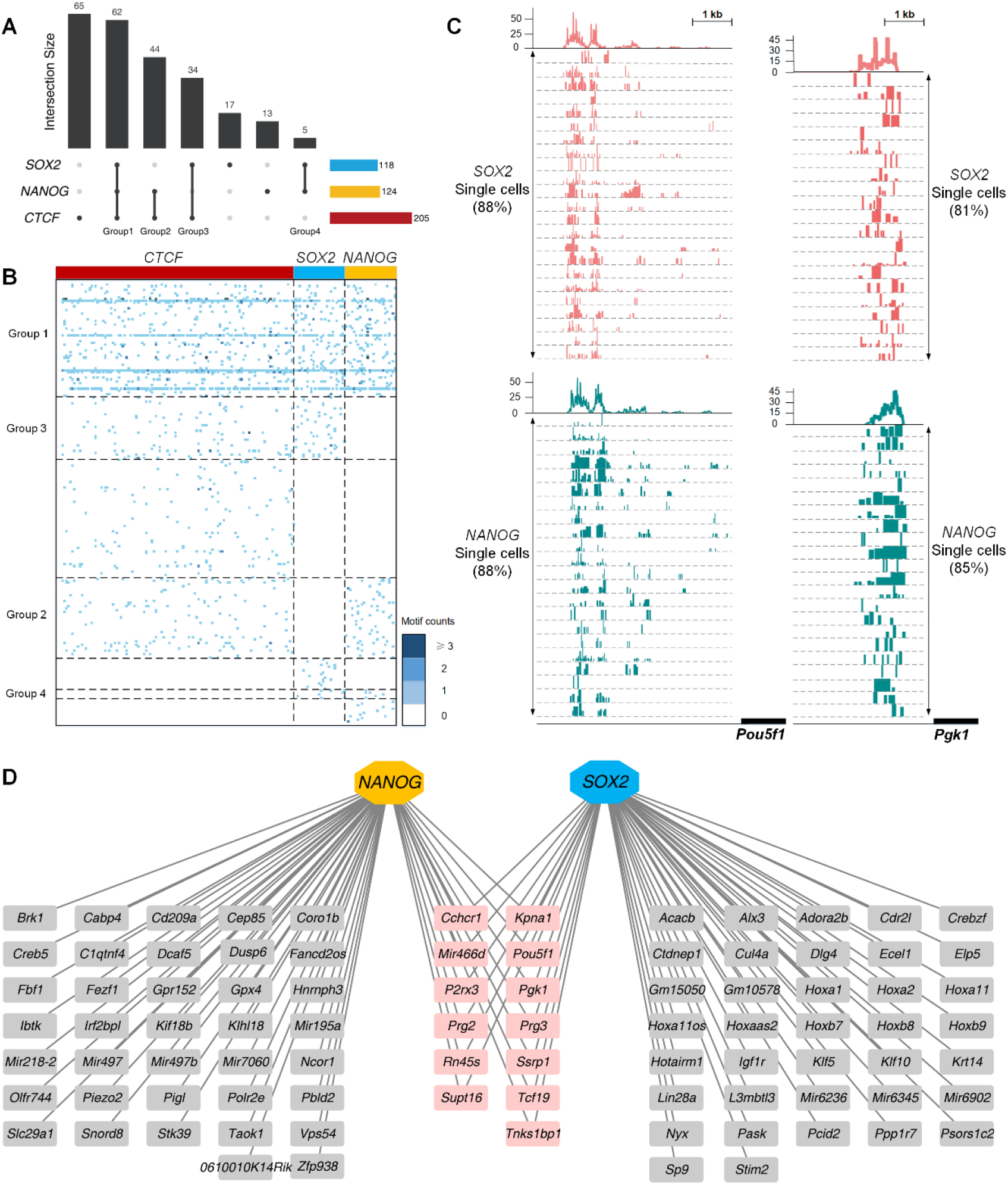
Single-cell CUT&RUN analysis. (A) An upset plot of the shared and unique motifs among CTCF, NANOG, and SOX2 datasets. (B) Motifs are found in each cell, and motifs identified are denoted as blue. (C) Co-enrichment of SOX2 and NANOG motifs in the upstream regulatory regions of genes Pou5f1 and Pgk1. (D) The regulatory network of SOX2 and NANOG is inferred from motif co-enrichment. The pink nodes represent the genes shared by NANOG and SOX2 with higher potentials to be co-regulated by SOX2 and NANOG.

With the observed motifs shared by different CUT&RUN data, we explored the relations between the motif co-enrichment and TF co-regulation. It has been reported that SOX2 and NANOG co-regulate the expression of Pou5f1 gene and maintain the pluripotency in human embryonic stem cells ^38^. We mapped the five motifs shared between SOX2 and NANOG data to the upstream 5kb region of *Pou5f1* using the mouse reference genome (i.e., GRCm38/mm10) and compared the mapping results with the peak region in the same region. As a result, we observed that 88% of SOX2 cells and 88% of NANOG cells had motif enrichment exactly in the same corresponding peak region, indicating that the motif co-enrichment of SOX2 and NANOG lead to the co-regulation of *Pou5f1* (**Fig. 5C**). Similarly, the motif co-enrichment of SOX2 and NANOG was also found in 3kb upstream regulatory region of *Pgk1*, so that we assume that *Pgk1* can also be co-regulated by SOX2 and NANOG.

To further investigate how SOX2 and NANOG co-regulate their downstream target genes, we constructed a gene regulatory network for each of the TF, respectively (**Fig. 5D**). The potential TF target genes were determined by matching identified motifs to the upstream promoter region via Cistrome-GO ^39^. We identified two groups of 37 genes that are solely regulated by SOX2 and NANOG, respectively, as well as 13 genes that are regulated by both TFs, including *Pgk1* and *Pou5f1*. Interestingly, both *Pou5f1* and *Pgk1* are reported as marker gene candidates to monitor embryo development stages^40^. Therefore, we reasoned that the rest of the 13 genes co-regulated by SOX2 and NANOG are also potential marker gene candidates to the mouse embryonic stem cells. Our result suggested that the existing DL methods are suitable for analyzing single-cell CUT&RUN data and can reveal potential TF co-regulatory patterns to potential marker gene candidates.

### 2.4 Develop the webserver to allow utilization of the DL models and accession of motif finding results

Despite the importance and high performance of the DL models in the above datasets, it is difficult to apply and utilize these models effectively in practical analyses. To address this issue, the Kipoi repository was recently accelerated community exchange and reuse of predictive models for genomics ^41^. However, programming skills and selecting suitable hardware are still required to apply DL models. Considering the results of our benchmark analysis, we developed a webserver to free the potential users from computational burden in using the DL models. This webserver allows users to browse comprehensive analysis results from this study in a one-step, user-friendly interface (https://bmbl.bmi.osumc.edu/deepmotif/).

Specifically, for the DESSO results, four major functionalities and visualizations were provided by the webserver (**Fig. 6A**). *(i)* Each of the identified sequences was available for comparisons (using TOMTOM) with existing databases, such as JASPAR^42^, HOCOMOCO^37^, Transfac^43^, and GOMo^44^ (**Fig. 6B**). A shape motif profile has ±50 bps flanking regions added using a bold orange curve (i.e. Minor Groove Width, Propeller Twist, Helix Twist, and Roll features computationally derived from DNA sequences by Monte Carlo simulation^45^). The two boundary curves of the blue region represented the upper and lower boundaries of the shape features in the corresponding motif instances. *(ii)* The motif instances and locations were identified (**Fig. 6C**). *(iii)* The line chart for the per-nucleotide vertebrate conservation and the corresponding mean value of motifs was available (**Fig. 6D-E**). Each figure has ±50 bps flanking regions added within the cell line. *(iv)* The enrichment score for the identified motifs was provided (**Fig. 6F**). The red curve indicated the enrichment score on its corresponding ChIP-Seq peaks. The vertical black lines indicated the presence of ChIP-Seq peaks that contain at least one TFBS.

**Fig. 6.**
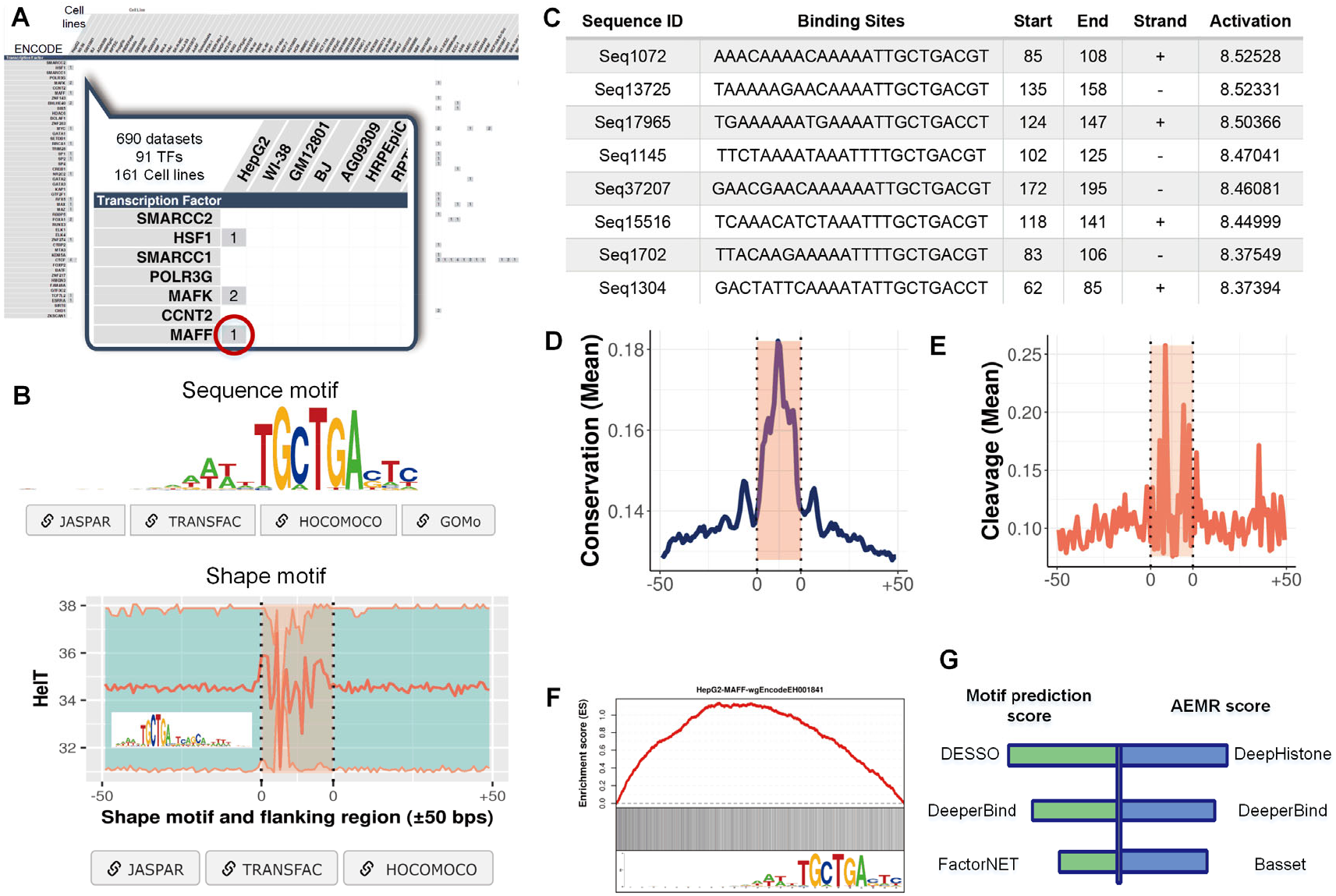
Overview of the webserver. (A) The matrix on the homepage, covering 690 ENCODE ChIP-Seq datasets with 161 cell lines and 91 TFs. Each entry indicates the number of datasets derived from a specific cell line and a specific TF. The TF MAFF dataset in the HepG2 cell line is highlighted and used as an example. (B) The identified MAFF sequence and shape motifs using DESSO, with links comparing their PWM to the existing motif databases. The shape motif profile has ±50 bps flanking regions added using a bold orange curve. The two boundary curves for the blue region represent the upper and lower boundaries of the shape features in the corresponding motif instances. (C) Snapshot of the MAFF TFBS table. (D) Line chart for the per-nucleotide vertebrate motif conservation and the ±50 bps flanking regions within the HepG2 cell line. (E) The line chart for the corresponding mean motif value and the ±50 bps flanking regions within the HepG2 cell line. (F) Enrichment score for an identified motif. The red curve indicates the enrichment score on its corresponding ChIP-Seq peaks. The motif logo is shown at the bottom. (G) Implemented tools in the webserver based on their superior performance in terms of the motif prediction and sequence classification scores.

Six deep learning frameworks were selected based on their overall scores (larger than 0.8) for each metric (**Fig. 6G**), and hence were implemented into the server to enable effective usage of these frameworks. They were DESSO, DeepBind, and FactorNET for motif prediction, and DeepHistone, DeeperBind, and Basset for sequence classification. The server allows users to submit their genomic sequencing data (e.g., ChIP-Seq and CUT&RUN) in the form of peak-lists through a submission page, which allows motif prediction or sequence classification. For each method, 690 pre-trained models from 91 TFs in 161 cell lines using ENCODE datasets are available to the user (**Supplementary Table S8**). The server currently supports two human genome versions (assembly GRCh38 and GRCh37). The output report, along with a unique job ID, will be generated and emailed to the user when the analysis is complete.

## 3. Discussion

Our benchmarking study includes a detailed evaluation of the performance of 20 DL tools on 1,043 genomic sequencing datasets, in terms of accuracy of motif finding, the performance of DNA/RNA sequence classification, algorithm scalability, and tool usability. Our quantitative evaluation provided a practical model and a set of optimal DL strategies for different datasets (ChIP-Seq, CLIP-Seq, cancer data, and single-cell data). Furthermore, a webserver was developed in our study, that enables the users to execute DL models on their own datasets, compare identified motifs from different datasets, and apply downstream analysis with a minimum of effort (e.g., finding genes near to the TFBS and analyzing enriched metabolic pathways). These new insights and computational infrastructures can significantly facilitate researchers in selecting the appropriate tools for their analyses related to motif finding and gene regulation.

Some practical challenges are also presented through our evaluation studies. First, the comparison results suggested that the existing models presented unstable performances for the 126 ChIP-Seq datasets obtained from nine cancer types. Second, the convolution kernel is used as the detector to scan the input sequence, however, the number and width of the convolution kernels have quite different settings in different tools. Specifically, the number of convolution kernels exceeds 100 in some DNA models, which will undoubtedly consume too much computing resources and produce redundant results. The width of the convolution kernel can be up to 26 in DNA models, while the width of the convolution kernel in RNA models is shorter. Third, other factors such as the presence of noisy data and class imbalance also need to be considered when choosing a DL model in practical motif finding.

Overall, the ‘good-yet-not-the-best’ methods can still provide a valuable contribution to motif finding when one takes advantage of novel algorithms, proposes a more scalable solution, or provides a unique insight in specific use cases. On the other hand, combining different tools in one analysis is always beneficial based on the observed complementarity of the evaluated DL models in our study. In the future, more emphasis needs to be placed on how to benchmark the existing methods and how to develop new methods with better robustness across different data types.

Furthermore, rather than measuring a single TF binding event in ChIP-Seq, a new technology, ATAC-Seq, can simultaneously detect hundreds of TF motif occurrences and has an average of 105,585 peaks from multiple cancer samples^46^. Considering the huge data scale and the fact that the majority of peaks are found in distal enhancers, inferring regulatory networks from ATAC-Seq data can be challenging. HINT-ATAC has been developed to identify TF footprints, showing less false positive TFBS results than the naïve peak-based method^47^. However, de novo motif discovery tools for ATAC-Seq data are lacking. Hence, the graph neural networks have been successfully used to predict various complex TF interactions and can potentially improve the reconstruction of regulatory networks from ATAC-Seq data.

## 4. Methods

### 4.1. Data collection

The 690 ChIP-Seq datasets were obtained from ENCODE, which covers 161 TFs and covered 91 human cell types for DNA models, and we selected the length of the sequence with 1,001bps as the input of these models. For the RNA models, we collected 55 CLIP-Seq datasets from reference paper ^23^ and used a fixed length of 501 bps as input. The Cistrome Data Browser was used to query the human cancer ChIP-Seq datasets and the selected 126 cancer ChIP-Seq of nine cancer types satisfy (*i*) the corresponding cell line should be a cancer cell, and (*ii*) the quality control standards of the peaks must be high. For single-cell datasets, 172 single-cell CUT&RUN datasets were retrieved from NCBI ^3^, including 120 CTCF, 26 NANOG, and 26 SOX2 cells. The cells contained TFBSs with lengths of 1 to 120 bps for each single-cell CUT&RUN dataset. All mined datasets, except the CLIP-Seq, only contained positive samples (peak sequences). Negative samples (e.g., randomly selected sequences) were generated following the strategy used in DESSO. In our experiments, each of the positive samples had a label of ‘1’, each of the negative samples had a label of ‘0’.

### 4.2. Data preprocessing

ChIP-Seq and CLIP-Seq datasets have been preprocessed in the original studies, the fixed length of sequences was directly available. As DL models require binary vector as input, each input sequences converted to an encoded matrix*M*= *L* × 4, *i.e. A* = [0, 0, 0, 1], *G* = [0, 1, 0, 0], *C* = [0, 0, 1, 0], *T* = [0, 0, 0, 1], where *L* is the length of input sequence^48^.

Regarding 126 cancer ChIP-Seq datasets and 172 single-cell CUT&RUN, the length of peaks in each dataset is different, so we pruned the original peak with a fixed length (101bps or 1001bps) by formula (1).

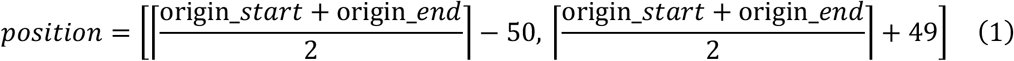

where origin_*start* is the start position of the original peak,origin_*end* is the end position of the original peak position, *position* is the position of the processed peak in the chromosome. After pruning, some redundant peaks may be generated and will be removed in the experiment, and the bedtools v2.21.0 was employed to acquire the pruned sequences^49^. Finally, the pruned sequences were encoded in the same way as ChIP-Seq and CLIP-Seq in the previous paragraph.

### 4.3. Model architecture

CNN, DNN, and DBN are classical DL algorithms, but only CNN was applied in each of 20 DL models. Hence, we divided the DNA models and the RNA models into two categories, *i.e.* CNN network and hybrid network. The CNN network category means models only contain convolution. The hybrid network category represents that models not only contain CNN but also RNN and/or DBN. Meanwhile, all the 20 DL models cover the eight component algorithms, which are convolution (upright pentagon), deconvolution (inverted pentagon), pooling (upright triangle), unpooling (inverted triangle), Dense (rectangle), Gate (grid circle), RNN (circle), DBN (rounded quadrilateral). Based on these components and their combinations, we grouped the 20 DL models into five types of architectures, which are conventional training, RNN training, multi-verse training, gate training, and parallel training (**Fig. 1C**).

### 4.4. DNA models and RNA models

All the 20 DL models are supervised learning methods^50^. According to different application scenarios, we divide the 20 DL models into two categories: the DNA models represent a type of model that can be applied to DNA sequences; and the RNA models represent a type of model that can be applied to the sequences. In practical applications, DeepBind, DanQ, DeeperBind, and Zeng’s models belong to the DNA model category have also been applied to RNA sequences, but none of the RNA models can be applied to DNA sequences.

### 4.5. Motif detection

Among the 20 DL models, ten of the 15 DNA models and three of the nine RNA models had the capability to find motifs. As described in Section 3.3 (Model architecture), all DL models can be divided into five categories, namely convolutional training, RNN training, multi-verse training, gate training, and parallel training. In summary, the DNA models identified motifs by scanning the one-hot encoding DNA sequences through convolutional kernels such as DeepBind, DanQ, and Basset. Three of the nine RNA models (iDeep, iDeepS, and iDeepE) were used to find motifs, and the convolution kernels also were used as motif detectors to find motifs (the same as the strategy used in the DNA models). The DL model was trained using ChIP-Seq or CLIP-Seq peaks, and each input was encoded as a matrix *M*_*L*×4_. Each convolution kernel of the trained model was seen as a detector to scan the input matrix *M*_*L*×4_, the final result *f*_*i*_ is given by formula (2).

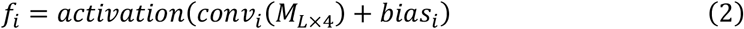

where, *f*_*i*_ is the convolutional result, which is a vector; *activation* represents activation function; *conv*_*i*_ is *i*^**th**^ convolution kernel; *bias*_*i*_ is the threshold value, the number of convolution kernels for each DL model is ranged in applications.

We selected a significant value of the vector *f*_*i*_ as the activation value to obtain the activated sequence with a predefined length. We hypothesized that the above motif detection way was the key to obtaining the best motif prediction scores. We obtained a set of aligned activated sequences and computed the position weight matrix (PWM)^51^. The number of columns of the matrix was four, and the number of rows of the matrix was the length of the convolution kernel. Each element in the matrix represented the number of base occurrences in the corresponding position. The PWM was aligned as the motif profiles and visualized using WebLogo ^52^. We identified the underlying TFs for the motifs by querying the HOCOMOCO v11 database through the TOMTOM v5.1.0 tool.

### 4.6. Model execution

Each execution of a model on a dataset was performed in a separate task, each task was allocated one CPU of Intel Xeon CPU E5-2683 v4 at 2.10 GHz and an NVIDIA Tesla P100 GPU. During the execution of a model on a dataset if the running time limit (>12h) was exceeded.

### 4.7. Evaluation workflow

All DL models were evaluated based on four metrics, including the AEMR metric that summed eight measurements for DNA/RNA sequence classification, the motif prediction score assesses the accuracy of motif finding, the algorithm scalability score assesses training time and efficiency, and the tool usability score assesses the quality of the tool implementation and usage. These four metrics were summed up as an overall score that was used to assess these DL models.

#### 4.7.1. AEMR score

We first calculate the precision, recall, F1_score, specificity, ACC, MCC, AUC, and PRC as shown below:

Precision is the ratio of the true predicted positive samples to all the predicted positive samples.

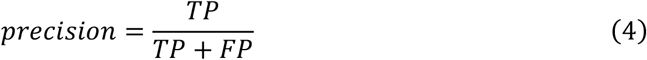

The recall represents the proportion of positive samples correctly predicted to the total positive samples.

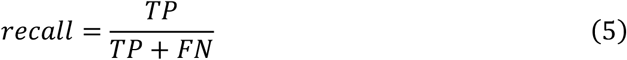

F1_score is the harmonic mean of precision and recall.

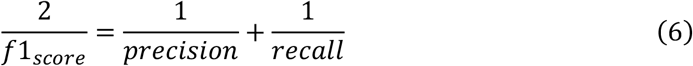

Specificity is the proportion of identified negative samples to all negative samples.

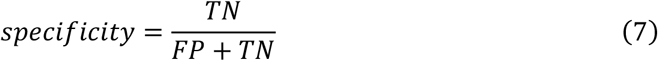

ACC is the proportion of correct prediction to the total sample.

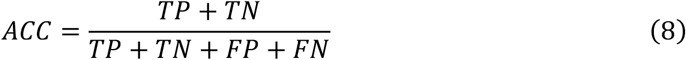

MCC is essentially the correlation coefficient between the observed and the predicted, it is a value between −1 and +1. The coefficient +1 means perfect prediction, 0 means no better than a random prediction, and −1 means discrepancy between prediction and observation.

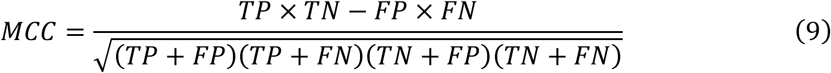

AUC is the area under the receiver operating characteristic curve, which is a value between 0 and 1. The closer AUC is to 1, the better the classification is.

PRC is the area under the precision-recall curve, which also is a value between 0 and 1. The difference between AUC and PRC is that when the positive and negative samples are unbalanced, PRC is more sensitive to experimental results.

Once all the eight scores are calculated, a radar chart can be generated that consists of eight equi-angular spokes with each spoke representing one of the scores defined above. The length of a spoke is proportional to the magnitude of the score for the data point relative to the maximum magnitude of the score across all data points. The AEMR score is the total area of the octagonal radar and is scaled to a score ranging from 0 to 1. The higher the AEMR score is, the better performance the tool has for sequence classification.

#### 4.7.2. Motif prediction score

The score assesses the accuracy of motif finding, which contains *P*-value, *E*-value, and *Q*-value of motif significance. The *P*-value represents the probability that a random motif, of the same width as a documented motif, would have an optimal alignment with a match score as good or better than the documented motif. Both *E*-value and *Q*-value was used as multi-testing correction, where the *E*-value indicates the expected number of false positives in the matches up to this point, and the *Q*-value is the minimal false discovery rate at which the observed similarity would be deemed significant. For each model, we calculate the median values of *P*-value, *E*-value, and *Q*-value of all found motifs, respectively, and further gave the motif prediction score as formula (10) and scaled from 0 to 1.

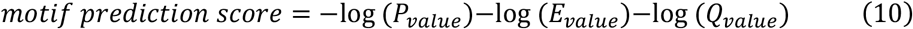

#### 4.7.3. Scalability score

To assess the scalability of each method, we started from four kinds of real datasets, each kind of dataset contained ten sub-datasets with a fixed number of peaks (10k, 20k, 30k, 40k). We run each model on each kind of dataset for maximally 12h. In order to determine the training time of each model, we started the timer after loading data and the package and stopped the timer after finishing the training process. And we recorded and normalized the averaged training time of each model on each kind of dataset, the scalability score was defined as the sum of the normalized training time in each kind of dataset.

#### 4.7.4. Usability score

To promote the improvement of current models and develop user-friendly new models, the usability of each model was quantified based on several existing model quality and programming guidelines (see details in **Supplementary Table 3**)^53^. The usability covered availability, behavior, code assurance, code quality, and documentation categories. Specifically, the availability category checked whether the packages and dependencies can be easily installed and whether the method is easily available and used. The code quality assessed the quality of the code both from a user perspective and a developer perspective. The code assurance category was frequently checked for code testing, continuous integration, and an active maintaining team. The documentation category checked the quality of the documentation, whether giving helpful documentation and clear tutorials. The behavior category evaluated the ease by which the method can be run, by looking for unexpected output files and messages and prior information. For each aspect, we also assigned a weight to the individual questions being investigated^53^, and each of the five categories was weighted equally for calculating the final score.

### 4.8. Webserver implementations

The server consolidated a range of web frameworks to provide user-friendly interactive visualizations of identified motifs. The back-end of the server was implemented in PHP for data queries and custom job submission. All data were stored and managed using a MySQL database. The entire web application was containerized with Docker and installed on a Red Hat Enterprise seven Linux system with 28-core Intel Xeon E5–2650 CPU and 64GB RAM.

### 4.9. Data availability

The assessable links and accession number of all datasets used in this study can be retrieved in **Supplementary Table S2**.

### 4.10. Code availability

All source codes of the 20 DL models can be found on GitHub, with links listed on https://bmbl.bmi.osumc.edu/deepmotif/. The evaluation code used in this study is available at https://github.com/OSU-BMBL/deepmotif-benchmark.

## Acknowledgments

This study was supported by the National Natural Science Foundation of China (Nos. 62072212, 61772227), the Development Project of Jilin Province of China (Nos. 20200401083GX, 2020C003), and Guangdong Key Project for Applied Fundamental Research (Grant 2018KZDXM076). This work was also supported by Jilin Provincial Key Laboratory of Big Data Intelligent Computing (No. 20180622002JC).

## Author information

### Contributions

Q.M designed the study. S.Z., Z.W., and C.W. performed the experiments and web portal establishment. Y.W., S.Z., D.X., and A.M prepared the manuscript. Y.W. and Q.M supervised the project.

