## Supplementary Figures for "Assessing deep learning algorithms in *cis*-regulatory motif finding based on genomic sequencing data"

**Contents**

**Figure S1**. Usability metrics assessment for the 20 DL tools.

**Figure S2**. Comparison of method ranks across different scoring metrics.

**Figure S3**. Eight metrics of DNA model on 126 cancer ChIP-seq datasets.

**Figure S4**. The average AEMR score across different DL models.

**Figure S5**. P-value of DNA model on 172 single-cell CUT&RUN datasets.

**Table S1.** All 20 benchmarked DL tools.

**Table S2.** Table S2. Datasets used for benchmark and cancer and single-cell investigation.

**Table S3.** Scoring sheet for assessing usability of DL models.

**Table S4.** AEMR scores, motif prediction scores, usability scores, scalability scores, and overall scores of DL models in this study.

**Table S5.** AEMR score of each DNA model, based on nine cancer types.

**Table S6.** Motifs found in each single-cell dataset.

**Table S7.** The shared motifs in four groups.

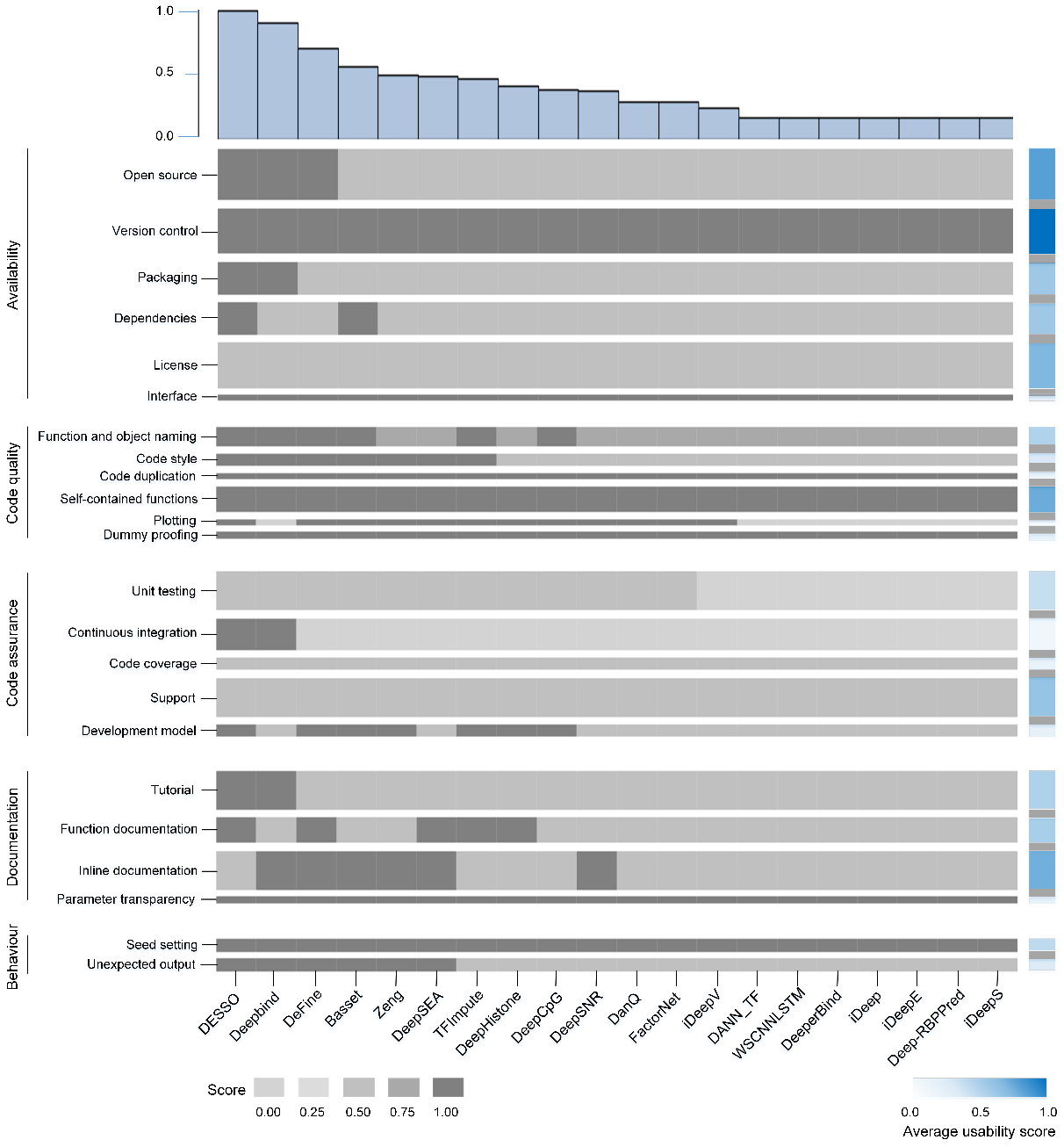

**Supplementary Fig. S1.** Usability metrics assessment for the 20 DL tools. Top: Average usability score for each model. Bottom left: metrics scores of each tool (Supplementary Table 3). Each category was weighted to equally contribute to the overall usability score. Bottom right: the average score of each quality control item.

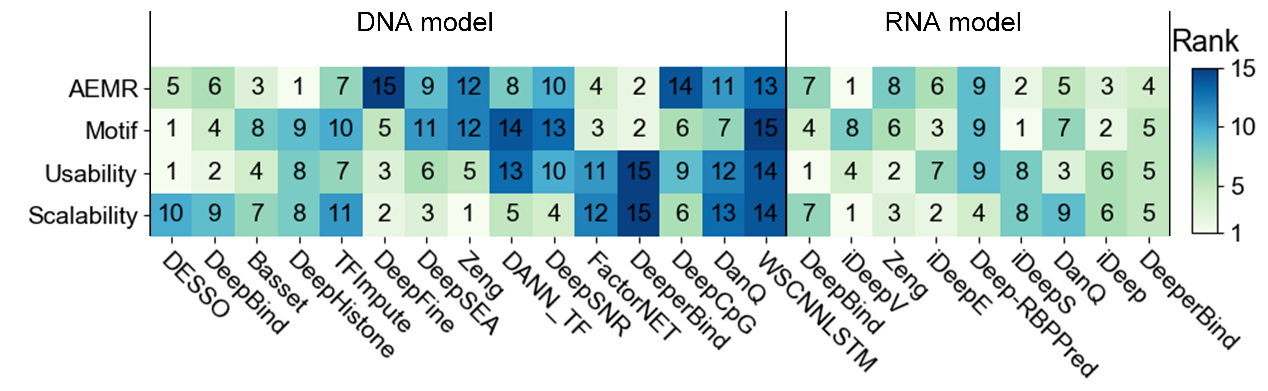

**Supplementary Fig. S2.** Ranks of the four evaluation criteria. Lower rank indicates better performance.

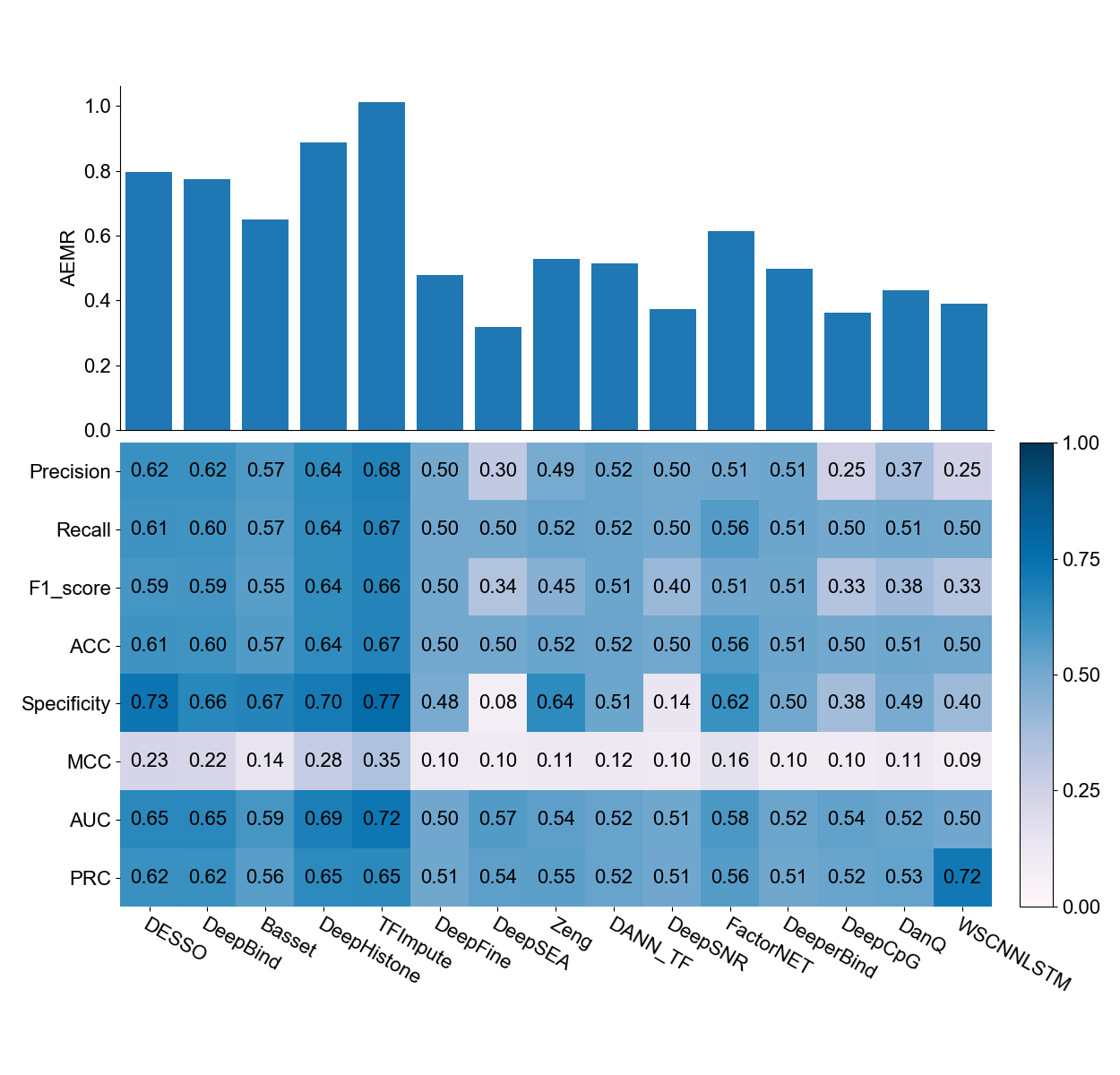

**Supplementary Fig. S3.** Eight metrics scores on 126 cancer ChIP-seq datasets.

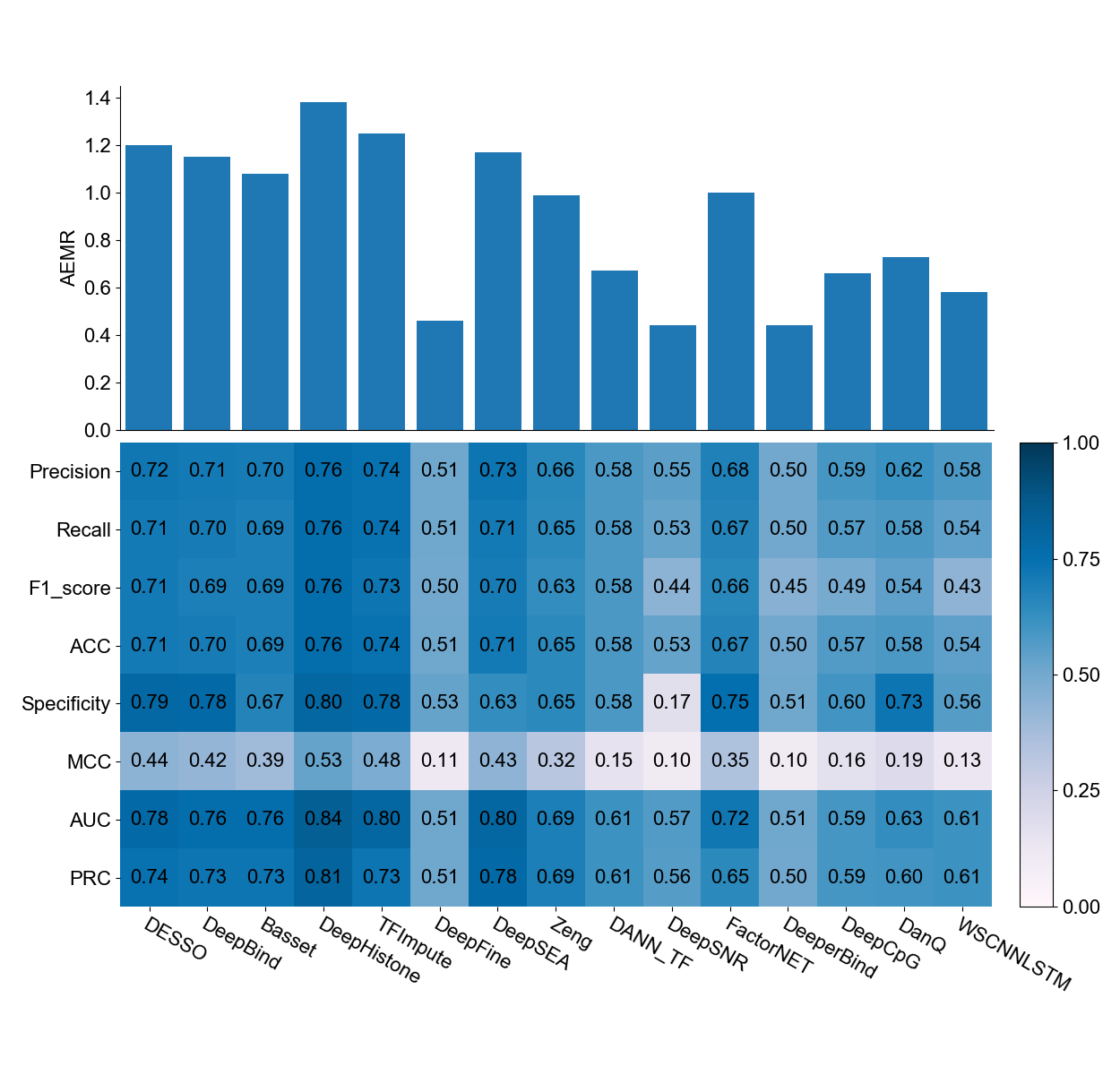

**Supplementary Fig. S4.** Eight metrics scores on 172 single-cell CUT&RUN datasets.

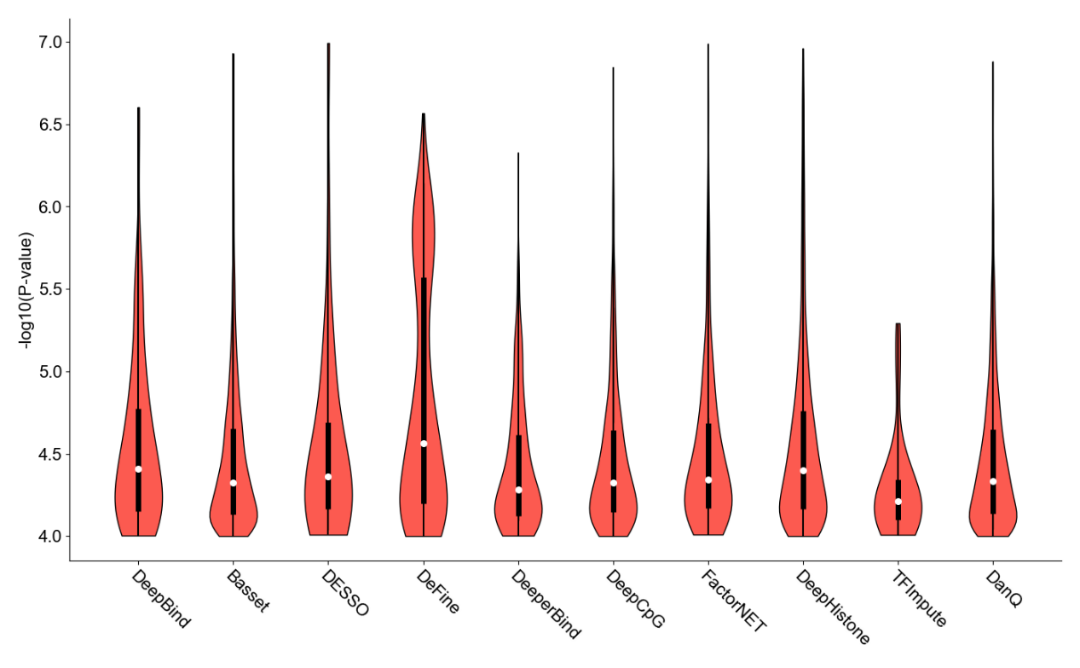

**Supplementary Fig. S5.** P-value of DNA model on 172 single-cell CUT&RUN datasets.
